# Parental origin of transgene determines recombination efficiency in GFAP-creERT2 mice

**DOI:** 10.1101/2025.02.27.640655

**Authors:** Karmen Mah, Adam Roszczyk, Ashley E. Frakes

**Affiliations:** Genetics and Biochemistry Branch, National Institute of Diabetes and Digestive and Kidney Diseases (NIDDK), National Institutes of Health, Bethesda, MD, USA; National Institute of Neurological Disorders and Stroke (NINDS), National Institutes of Health, Bethesda, MD, USA

## Abstract

The Cre-loxP system is a powerful tool for spatial and temporal genetic manipulations. However, the system is prone to several limitations and caveats with respect to variable expression and recombination in unintended cell types. Using one of the most widely used astrocyte Cre lines, hGFAP-creERT2 (GFAP-cre/ERT2)505Fmv/J), we found that parental origin of the hGFAP-creERT2 transgene is a determinant of recombination efficiency. Recombination was robust in animals with paternally inherited hGFAP-creERT2. However, animals with maternally inherited Cre exhibited little to no recombination. This result was recapitulated using female hGFAP-creERT2 mice procured directly from Jackson Laboratory. We did not observe transgenerational suppression of Cre recombination in maternally inherited hGFAP-CreERT2. These data highlight the need for careful planning and documentation of breeding schemes when working with Cre mice.

## Introduction

Astrocytes are one of the largest cell populations in the central nervous system (CNS) and reside in all brain and spinal cord regions. They perform many functions including supporting neuronal development and survival, buffering pH and ion concentrations, and forming the blood brain barrier (Khakh & Deneen, 2019). Astrocytes also contribute to pathological conditions such as neurodevelopmental and neurodegenerative diseases. Therefore, it is essential to have specific and reliable tools to genetically manipulate astrocytes to reveal the mechanisms that regulate astrocyte function in health and disease(Yu et al., 2020).

The Cre-loxP system has been the most widely used approach for spatial and temporal genetic manipulations in mice. Cre recombinase recognizes 34 base pair loxP sites which can be leveraged to delete genes by flanking the DNA of interest with loxP sites (floxed alleles). The development of tissue-specific Cre driver lines and floxed alleles in mice, has enabled genetic manipulation for many cell types in the CNS. However, several caveats to the Cre-loxP system have been reported such as Cre expression in unintended cell types, germline recombination, and mosaic or inconsistent recombination(Luo et al., 2020). To avoid caveats associated with Cre expression during development, creERT2 lines were developed to allow for temporal control of Cre activity. CreERT2 is a fusion protein of Cre recombinase and the mutated human estrogen receptor and is activated by the administration of tamoxifen or its metabolite, 4-hydroxytamoxifen (4-HT) (Feil et al., 1997).

Many astrocyte CreERT2 drivers have been developed with varying specificity and efficiency such as human glial acidic fibrillary protein (hGFAP)-CreERT2, glutamate aspartate transporter (Glast)-CreERT2, connexin 30 (Cx30)-CreERT2, fibroblast growth factor receptor 3 (Fgfr3)-CreERT2 and aldehyde dehydrogenase 1 family member L1 (Aldh1l1)-CreERT2 (Casper et al., 2007; Ganat et al., 2006; Hu et al., 2020; Mori et al., 2006; Srinivasan et al., 2016; Winchenbach et al., 2016; Young et al., 2010). Aldh1L1-creERT2 targets the most astrocytes with high specificity in the brain. However, Cre activity is also detectable in peripheral tissues in Aldh1L1-CreERT2 mice including the liver, lungs, heart, spleen, intestine, and muscle (Hu et al., 2020). Glast- and Cx30-CreERT2 activity is also detectable in in two morphologically distinct lung cells and in the walls of blood vessels (Hu et al., 2020).

To avoid Cre recombination in peripheral tissues for our studies, we selected the hGFAP-CreERT2 mouse strain (B6.Cg-Tg(GFAP-cre/ERT2)505Fmv/J), one of the most widely used astrocyte Cre drivers. After observing variability in recombination efficiency, we systematically evaluated Cre recombination efficiency based on the parental origin of the hGFAP-CreERT2 allele. We bred either male or female hGFAP-CreERT2(+) animals to loxP-stop-loxP-TdTomato Cre reporter animals (LSL-TdTomato). After tamoxifen administration to activate CreERT2, the floxed stop cassette is removed in Cre-positive cells, resulting in TdTomato expression as a marker for Cre activity. We found that experimental animals with maternally inherited Cre exhibited little to no TdTomato-positive cells, indicating no recombination. This observation was recapitulated using female hGFAP-CreERT2 animals procured directly from The Jackson Laboratory. However, recombination was robust in animals with paternally inherited hGFAP-CreERT2. We did not observe transgenerational suppression of Cre recombination in maternally inherited hGFAP-CreERT2. These data highlight the importance of careful planning and documentation of breeding schemes when working with Cre mice.

## Results

### Parental origin of GFAP-CreERT2 transgene influences recombination efficiency

To determine if hGFAP-CreERT2 recombination efficiency is influenced by the parental source of Cre, we bred either male or female hGFAP-CreERT2(+) animals to lox-stop-lox-TdTomato Cre reporter animals (LSL-TdTomato). We injected hGFAP-CreERT2(+);LSL-TdTomato(Tg/wt) progeny with tamoxifen via intraperitoneal (ip) injection for 5 days at 5-6 weeks of age. We harvested brain tissues 3 weeks after the last tamoxifen injection and assessed TdTomato fluorescence (Fig 1A). We scored brain regions for TdTomato fluorescence as high, medium, low, or none corresponding to >70%, 40-70%, <40%, and 0% cells that are TdTomato+, respectively (Fig 1B). As expected, TdTomato fluorescence colocalized with astrocyte markers S100β or Sox9 indicating specific Cre recombination in astrocytes (Fig 1B, merge).

**Figure 1:**
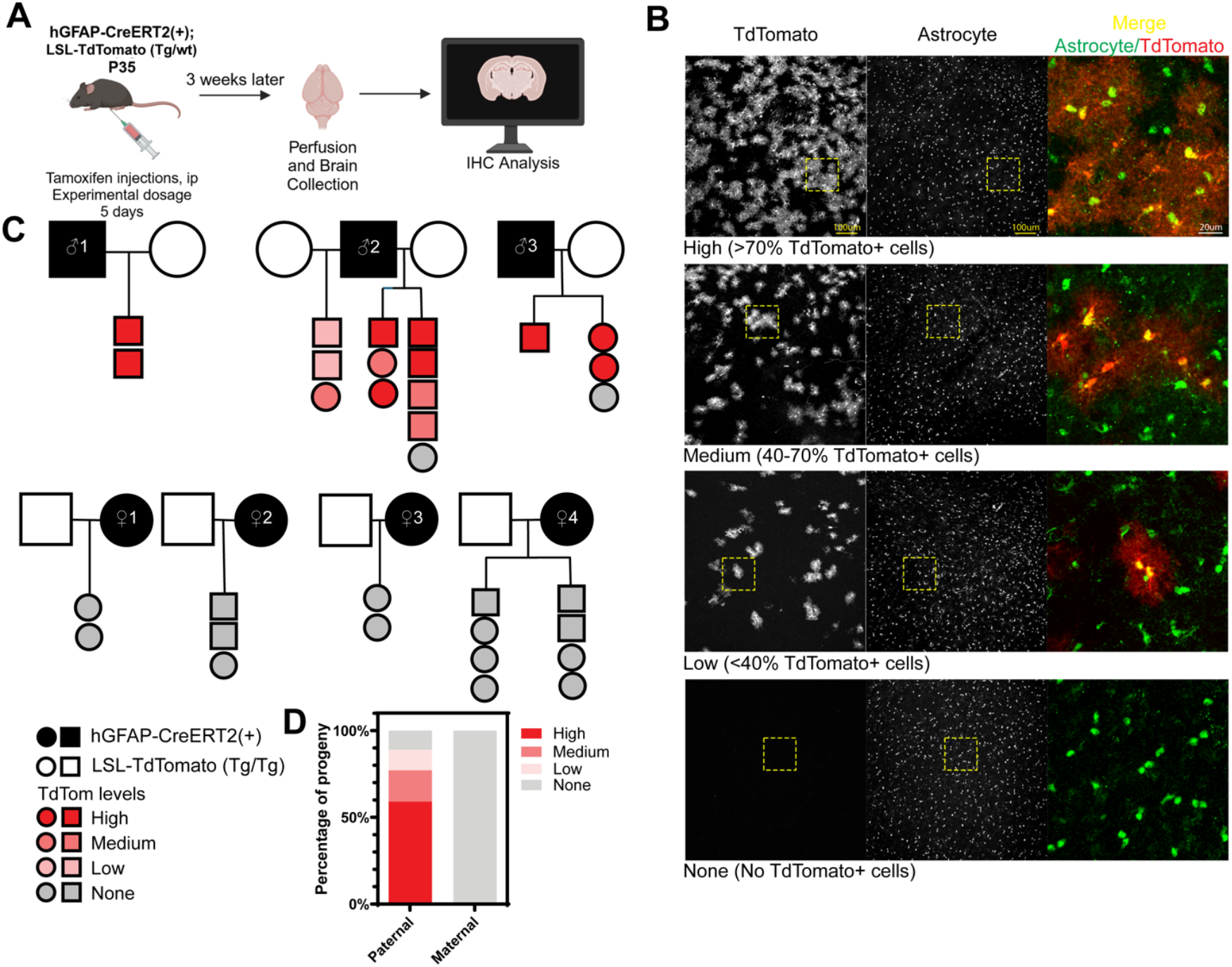
Parental origin of GFAP-CreERT2 transgene influences recombination efficiency. **A**. Schematic of the experiment. hGFAP-CreERT2(+);LSL-TdTomato(Tg/wt) animals were injected with tamoxifen via ip for 5 days. 3 weeks after the last injection, animals were perfused and prepared for immunohistochemistry to assess TdTomato fluorescence. **B**. Characterization of TdTomato fluorescence ratings. Representative hypothalamus micrographs of animals with high (>70% TdTomato+ cells in the brain), medium (40-70% TdTomato+ cells in the brain), low (<40% TdTomato+ cells in the brain) and none (no TdTomato+ cells in the brain). From left to right: TdTomato, astrocyte marker (S100b or human Sox9), and zoomed in magnification of the yellow dashed box showing the merge from TdTomato and astrocyte marker. Scale bars are 100 μm and 20 μm. **C**. Pedigree of experimental animals. Squares=males, circles=females. Black-filled squares or circles indicate hGFAP-CreERT2(+) carriers. Unfilled squares and circles indicate LSL-TdTomato (Tg/Tg) animals. Only hGFAP-CreERT2(+);LSL-TdTomato(Tg/wt) progeny are shown in the pedigree. Animals are colored with shades of red to grey to indicate the TdTomato fluorescence rating of the animal. **D**. Percentage of progeny grouped by the parent-of-origin of hGFAP-CreERT2 allele. About 90% of animals inheriting the hGFAP-CreERT2 allele paternally express TdTomato, while 0% of animals inheriting the allele maternally express TdTomato. The bar is colored red to grey representing the TdTomato fluorescence rating of the animals. N = 17 (paternal), N=15 (maternal).

Next, we analyzed the pedigrees and found that every animal that inherited CreERT2 from the maternal parent failed to undergo recombination and induce TdTomato reporter gene expression post tamoxifen administration. Across four breeding pairs with hGFAP-CreERT2(+) females, no progeny (n=15) exhibited TdTomato+ cells in the brain (Fig 1D). On the contrary, when the hGFAP-CreERT2 (+) allele was inherited paternally, we found that 17 of 19 animals had TdTomato+ cells. We observed some variability in the extent of Cre-mediated recombination in progeny with paternally-derived Cre. For example, Male #1 produced all offspring that had a high TdTomato fluorescence rating (Fig 1C). Males #2 and #3 each had a single female mouse with no TdTomato expression. Male #2 produced offspring that exhibited varying levels of TdTomato fluorescence ratings across multiple litters produced with two different LSL-TdTomato (Tg/Tg) females. These data suggest that parental source of CreERT2 is a determinant in whether or not hGFAP-creERT2 can drive recombination.

### Mice exhibit suppression of Cre activity when hGFAP-creERT2 is inherited from female mice directly from Jackson Laboratory

To determine if the transgenic inheritance effect was specific to our colony, we obtained hemizygous hGFAP-CreERT2(+) and LSL-TdTomato(Tg/Tg) males directly from The Jackson Laboratory to use as breeders to generate experimental animals (Fig 2A). We injected progeny with tamoxifen via intraperitoneal injection for 5 days and then assessed for TdTomato fluorescence 3 weeks after the last injection. Again, we observed a striking lack of recombination in hGFAP-CreERT2(+);LSL-TdTomato(Tg/wt) progeny that inherited Cre from the maternal parent. Female #5 and #7 produced no progeny with TdTomato+ cells. Female #6 produced two litters where only 1 out of 5 total animals exhibited low levels of TdTomato fluorescence. Thus, only 1 out of 9 progeny from breeder animals procured directly from The Jackson Laboratory had any TdTomato+ cells (Fig 2B). These data demonstrate that failure of recombination by maternally derived CreERT2 is not restricted to animals that were bred in-house and might reflect a phenotype pertaining to a broader population of animals.

**Figure 2:**
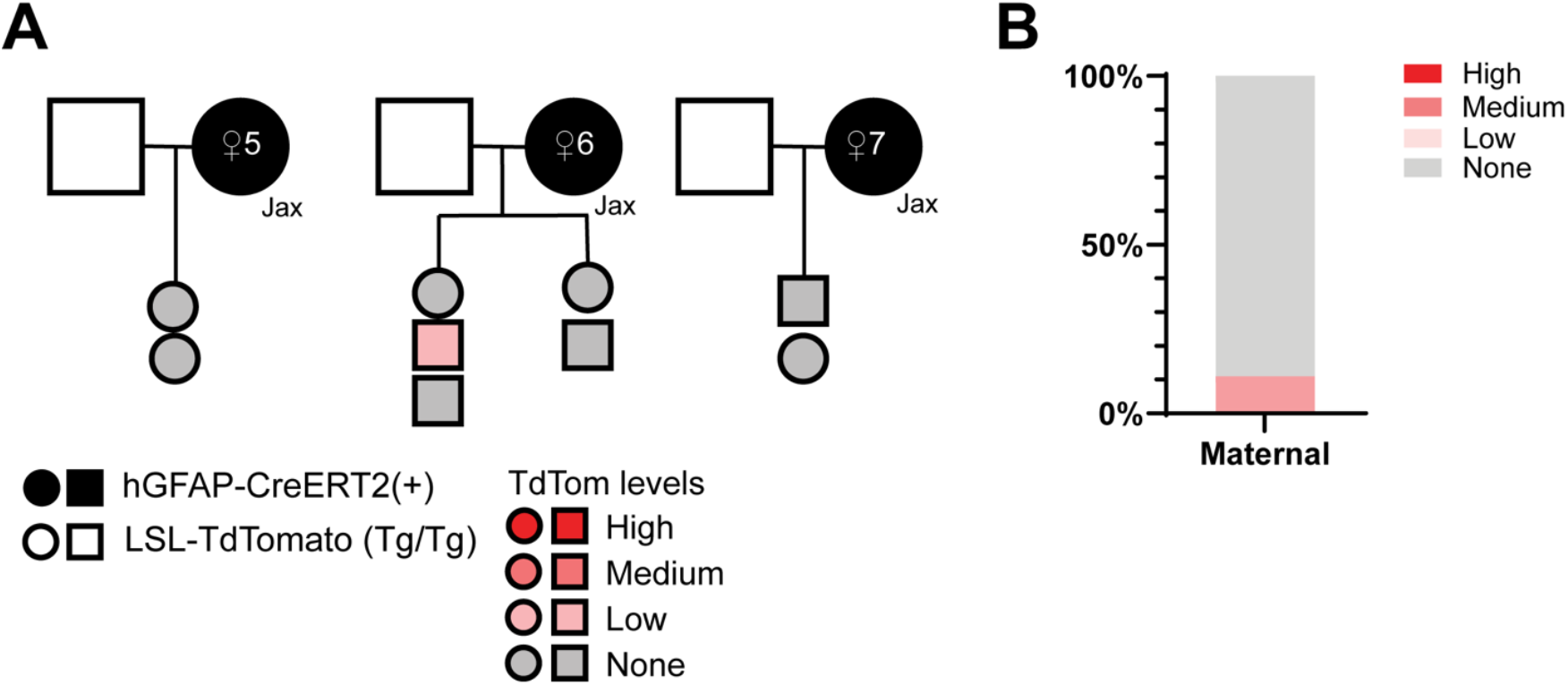
Mice exhibit suppression of Cre activity when hGFAP-creERT2 is inherited from female mice directly from Jackson Laboratory. **A**. Pedigree of hemizygous female hGFAP-CreERT2(+) animals procured from The Jackson Laboratory. Squares=males, circles=females. Unfilled squares represent LSL-TdTomato(Tg/Tg) males, black-filled circles indicate hGFAP-CreERT2(+) females from The Jackson Laboratory. Progeny are filled in with shades of red to grey indicating the TdTomato fluorescence rating, from high to none. **B**. Percentage of progeny originating from female carriers from The Jackson Laboratory and their fluorescence rating. 10% of animals had low TdTomato fluorescent rating, while the remaining 90% had no TdTomato+ cell. N = 9.

### Suppression of Cre recombinase activity by maternally inherited hGFAP-creERT2 is not transgenerational

As we observed no TdTomato fluorescence in animals inheriting the hGFAP-CreERT2 allele maternally, we sought to determine if this lack of Cre recombinase activity persists in animals transgenerationally. To test for transgenerational suppression of maternal hGFAP-CreERT2, we set-up breeding pairs consisting of a female hGFAP-CreERT2(+) animal (Fig 3B, female #10) and C57/Bl6 male or vice versa (Fig 3A, male #4). Resulting CreERT2(+) mice (both male and female) were mated with LSL-TdTomato (Tg/Tg) animals to produce a second generation of progeny in which recombination efficiency was evaluated. As expected, progeny with paternally inherited hGFAP-CreERT2 from two generations exhibit a high percentage of TdTomato+ cells (Male #4 and #5, Fig 3A). Progeny with maternally inherited hGFAP-CreERT2 show no TdTomato+ cells, indicated no Cre recombination, even when the female hGFAP-CreERT2(+) breeders inherited Cre paternally (Fig 3A, female #8 and #9). Interestingly, when a male hGFAP-CreERT2(+) breeder inherited Cre maternally, its progeny exhibited robust TdTomato fluorescence (Fig 3B, male #6). However, female littermates that were used as hGFAP-CreERT2(+) breeders (female #11 and #12) generated progeny that do not exhibit TdTomato fluorescence, indicating no Cre recombination (Fig 3B). These results demonstrate that the capacity for Cre recombination is determined by the sex of the immediate parent-of-origin of the GFAP-CreERT2 allele and is not transmitted transgenerationally.

**Figure 3:**
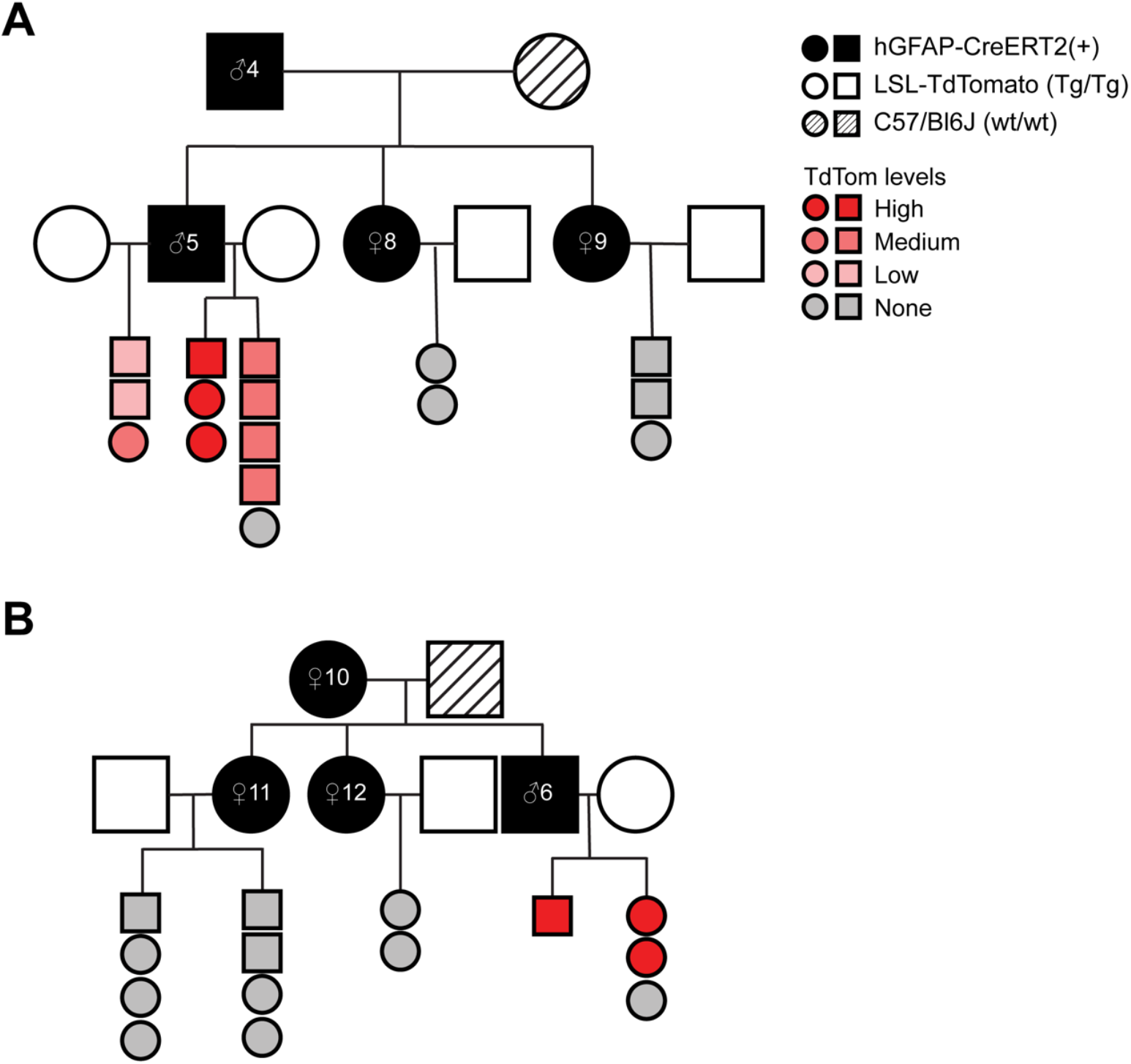
Suppression of Cre recombinase activity by maternally inherited hGFAP-creERT2 is not transgenerational. **A**. Pedigree of animals inheriting hGFAP-CreERT2 from a parental carrier. Squares indicate male animals, while circles are females. Black-filled squares or circles indicate hGFAP-CreERT2(+) carriers. Hatched-filled circle indicate C57/Bl6 female. Unfilled squares and circles indicate LSL-TdTomato (Tg/Tg) animals. Only hGFAP-CreERT2(+);LSL-TdTomato(Tg/wt) progeny are shown in the pedigree. Animals are colored with shades of red to grey to indicate the TdTomato fluorescence rating of the animal. **B**. Pedigree of animals inheriting hGFAP-CreERT2 from a maternal carrier.

## Discussion

The hGFAP-CreERT2 mouse strain is one of the most widely used Cre drivers to temporally control gene expression in astrocytes. After observing variability in recombination efficiency, we sought to systematically evaluate Cre recombination efficiency based on the parental origin of the hGFAP-CreERT2 allele. We crossed hGFAP-CreERT2 animals to the LSL-TdTomato Cre reporter mouse strain that expresses the TdTomato fluorescent protein following the excision of a loxP-flanked STOP cassette. Thus, the number of TdTomato-positive cells represents the recombination frequency in this mouse strain. We found that experimental animals inheriting CreERT2 maternally exhibited little to no TdTomato-positive cells indicating no recombination. This observation was recapitulated using female hGFAP-CreERT2 animals procured directly from The Jackson Laboratory. Interestingly, we do not observe transgenerational suppression of Cre recombination of maternally inherited hGFAP-CreERT2.

While we observed silencing of maternally inherited hGFAP-CreERT2 in both our in-house colony and animals directly acquired from Jackson Labs, it is possible that not all colonies of hGFAP-CreERT2 mice demonstrate this phenotype. We obtained our first hGFAP-creERT2 breeders from Jackson Laboratory in 2022 when we were starting our mouse colony. Additional breeders used in Fig. 2 were obtained in 2024. We cannot conclude whether hGFAP-CreERT2 animals distributed from The Jackson Laboratory prior to 2022 exhibit maternal suppression of Cre recombination. Therefore, we urge all investigators using this strain to test Cre recombination efficiency from male and female hGFAP-CreERT2 breeders in their colony. Furthermore, different loci can exhibit different sensitivities to Cre recombination, so testing recombination efficiency for all floxed alleles is encouraged (Becher et al., 2018).

Our results are not the first to demonstrate that parental inheritance of the Cre allele can influence recombination efficiency. For example, two germ cell-specific Cre drivers, Ella-Cre and Vasa-Cre, are more efficient if inherited maternally (Gallardo et al., 2007; Heffner et al., 2012). Many nervous system-specific Cre drivers also exhibit maternal or paternal germline recombination (Luo et al., 2020). These studies demonstrate the need for careful documentation and reporting of breeding schemes and sexes of experimental animals used.

In addition to Cre recombinase, other transgenes exhibit phenotypic differences based on parental origin. For example, in the widely used 5XFAD mouse model of Alzheimer’s disease, 5XFAD mice exhibit a more severe phenotype if the transgene is paternally inherited (Sasmita et al., 2025). Maternal inheritance lowers expression of the Thy1.2-promoter driven transgene which results in a lower amyloid plaque burden. While the mechanism contributing to this difference remain unknown, it was proposed that genomic imprinting may be involved. Genomic imprinting controls whether the maternal or paternal copy of a gene is expressed by epigenetic silencing of one allele. Therefore, it is possible that epigenetic modifications in the female germline reduce expression of the 5XFAD transgene.

Parent-of-origin effects have also been observed in more tractable model organisms such as *Caenorhabditis elegans* and *Caenorhabditis tropicalis* allowing for further mechanistic insight. Transgene silencing was discovered in a *C. elegans* strain that expresses single copy tandem transgenes of mCherry and GFP in the germline. Only animals that inherit the transgene through male sperm exhibit silencing, coining the term “mating-induced silencing”. This reliable and reproducible silencing was observed in >1500 animals generated from over 140 independent crosses. This silencing requires components of the small RNA pathway and can persist for several generations (Devanapally et al., 2021). In a parallel line of questioning, research in *C. tropicalis* found that the *slow-1/grow-1* toxin-antidote element was inactive when inherited paternally (Pliota et al., 2024). This phenomenon was attributed to the transcriptional repression of the element by PIWI-interacting RNA host defense pathway. These studies highlight the power of simple model systems when dissecting complex mechanisms that give rise to phenotypic differences based on inheritance.

Recent advancements in adeno-associated viral vector (AAV) design and delivery provide alternative tools to achieve spatial and temporal gene recombination in astrocytes. AAVs are safe, exhibit little long-term toxicity and specific serotypes show a high tropism for the CNS such as AAV-9 and AAV-PHP.eB (Chan et al., 2017; Foust et al., 2009). To improve astrocyte specificity of viral constructs, a multiplexed cassette of six microRNA-targeting sequences can be added to induce transgene degradation in neurons (Gleichman et al., 2023). This toolbox of vectors provides a highly specific method to manipulate genes in astrocytes, offering an alternative to traditional CreERT2 mouse models.

Taken together, our data demonstrate that hGFAP-CreERT2-mediated recombination of LSL-TdTomato only occurs when paternally inherited. Although the mechanism giving rise to suppression of Cre recombination in maternally inherited hGFAP-CreERT2 is unknown, our work highlights the importance of careful documentation of breeding schemes and characterization of Cre drivers. We have submitted our observations to the The Jackson Laboratory Cre characterization resource (https://www.creportal.org/) (Heffner et al., 2012).

## Methods

### Animals

Mouse husbandry and experimental procedures were approved by the National Institutes of Health Animal Care and Use Committee. Mice were housed in 22–24°C with a 12-hour light/12-hour dark cycle. Mice were fed with *ad libitum* standard chow (NIH-07; LabDiet Cat#5018) and water. The following mouse strains mice were purchased from The Jackson Laboratory: GFAP-creERT2 (B6.Cg-Tg(GFAP-cre/ERT2)505Fmv/J, Strain no: 012849) and the LSL-TdTomato Ai14 Cre reporter line (B6.Cg-*Gt(ROSA)26Sor*^*tm14(CAG-tdTomato)Hze*^/J, Strain no. 007914), and C57/Bl6J animals (Strain no: 000664). All mice were genotyped a minimum of two times by Transnetyx and in-house PCR-based methods.

### Tamoxifen treatment

Tamoxifen (T5648; Sigma Aldrich) is dissolved in Corn Oil (CO136; Spectrum Chemical) at 20mg/ml and filter sterilized. Animals were injected at 5-6 weeks old at a dose of 150mg/kg intraperitoneally (i.p.) for 5 days. Doses were skipped if animals lost >10% of body weight at start of injection series.

### Immunohistochemistry

Mice were anesthetized with Avertin (250mg/kg) and processed for transcardiac perfusion with saline (PBS) and 4% paraformaldehyde (PFA). Mouse brains were harvested and postfixed in 4% PFA at least overnight. Brains were embedded in 2% agarose before 40um coronal sections were cut on a vibratome (Leica VT1000 S). Sections were collected and blocked with blocking solution (10% Donkey Serum in TBS with 0.3% Triton X-100), before incubating in primary antibody diluted in blocking solution (Rabbit anti-S100b (Abcam ab52642), 1:5000; Goat anti-Sox9 (R&D AF3075) 1:250; Rb anti-dsRed (TakaraBio 632496) 1:500; Mouse anti-NeuN (EMD Millipore MAB377) 1:400) for 3 nights rocking at 4°C. Tissues were washed in TBS with 0.3% Triton X-100 for 3 times before incubating with the appropriate secondary antibodies diluted in blocking solution (all secondaries were used at 1:500; Jackson Immunoresearch, 711-545-152, 711-295-152, 715-605-150, 705-545-003) diluted in blocking solution for 2 hours at room temperature. Tissues were washed in TBS with 0.3% Triton X-100 3 times. Tissues were then mounted on slides and coverslipped with Vectashield Antifade Mounting Media (VectorLabs). Images were captured on a Leica Thunder Imager.

## Author contributions

KM performed injections, KM and AR managed breeding, genotyping, performed tissue processing and data collection. AEF and KM wrote the manuscript.

## References

Becher, B., Waisman, A., & Lu, L.-F. (2018). Conditional Gene-Targeting in Mice: Problems and Solutions. Immunity, 48(5), 835–836. 10.1016/j.immuni.2018.05.002

Casper, K. B., Jones, K., & McCarthy, K. D. (2007). Characterization of astrocyte-specific conditional knockouts. Genesis, 45(5), 292–299. 10.1002/dvg.20287

Chan, K. Y., Jang, M. J., Yoo, B. B., Greenbaum, A., Ravi, N., Wu, W.-L., Sánchez-Guardado, L., Lois, C., Mazmanian, S. K., Deverman, B. E., & Gradinaru, V. (2017). Engineered AAVs for efficient noninvasive gene delivery to the central and peripheral nervous systems. Nature Neuroscience, 20(8), 1172–1179. 10.1038/nn.4593

Devanapally, S., Raman, P., Chey, M., Allgood, S., Ettefa, F., Diop, M., Lin, Y., Cho, Y. E., & Jose, A. M. (2021). Mating can initiate stable RNA silencing that overcomes epigenetic recovery. Nat Commun, 12(1), 4239. 10.1038/s41467-021-24053-4

Feil, R., Wagner, J., Metzger, D., & Chambon, P. (1997). Regulation of Cre recombinase activity by mutated estrogen receptor ligand-binding domains. Biochem Biophys Res Commun, 237(3), 752–757. 10.1006/bbrc.1997.7124

Foust, K. D., Nurre, E., Montgomery, C. L., Hernandez, A., Chan, C. M., & Kaspar, B. K. (2009). Intravascular AAV9 preferentially targets neonatal neurons and adult astrocytes. Nat Biotechnol, 27(1), 59–65. 10.1038/nbt.1515

Gallardo, T., Shirley, L., John, G. B., & Castrillon, D. H. (2007). Generation of a germ cell-specific mouse transgenic Cre line, Vasa-Cre. Genesis, 45(6), 413–417. 10.1002/dvg.20310

Ganat, Y. M., Silbereis, J., Cave, C., Ngu, H., Anderson, G. M., Ohkubo, Y., Ment, L. R., & Vaccarino, F. M. (2006). Early postnatal astroglial cells produce multilineage precursors and neural stem cells in vivo. J Neurosci, 26(33), 8609–8621. 10.1523/jneurosci.2532-06.2006

Gleichman, A. J., Kawaguchi, R., Sofroniew, M. V., & Carmichael, S. T. (2023). A toolbox of astrocyte-specific, serotype-independent adeno-associated viral vectors using microRNA targeting sequences. Nature Communications, 14(1), 7426. 10.1038/s41467-023-42746-w

Heffner, C. S., Herbert Pratt, C., Babiuk, R. P., Sharma, Y., Rockwood, S. F., Donahue, L. R., Eppig, J. T., & Murray, S. A. (2012). Supporting conditional mouse mutagenesis with a comprehensive cre characterization resource. Nat Commun, 3, 1218. 10.1038/ncomms2186

Hu, N. Y., Chen, Y. T., Wang, Q., Jie, W., Liu, Y. S., You, Q. L., Li, Z. L., Li, X. W., Reibel, S., Pfrieger, F. W., Yang, J. M., & Gao, T. M. (2020). Expression Patterns of Inducible Cre Recombinase Driven by Differential Astrocyte-Specific Promoters in Transgenic Mouse Lines. Neurosci Bull, 36(5), 530–544. 10.1007/s12264-019-00451-z

Khakh, B. S., & Deneen, B. (2019). The Emerging Nature of Astrocyte Diversity. Annu Rev Neurosci, 42, 187–207. 10.1146/annurev-neuro-070918-050443

Luo, L., Ambrozkiewicz, M. C., Benseler, F., Chen, C., Dumontier, E., Falkner, S., Furlanis, E., Gomez, A. M., Hoshina, N., Huang, W. H., Hutchison, M. A., Itoh-Maruoka, Y., Lavery, L. A., Li, W., Maruo, T., Motohashi, J., Pai, E. L., Pelkey, K. A., Pereira, A., … Craig, A. M. (2020). Optimizing Nervous System-Specific Gene Targeting with Cre Driver Lines: Prevalence of Germline Recombination and Influencing Factors. Neuron, 106(1), 37-65.e35. 10.1016/j.neuron.2020.01.008

Mori, T., Tanaka, K., Buffo, A., Wurst, W., Kuhn, R., & Gotz, M. (2006). Inducible gene deletion in astroglia and radial glia--a valuable tool for functional and lineage analysis. Glia, 54(1), 21–34. 10.1002/glia.20350

Pliota, P., Marvanova, H., Koreshova, A., Kaufman, Y., Tikanova, P., Krogull, D., Hagmüller, A., Widen, S. A., Handler, D., Gokcezade, J., Duchek, P., Brennecke, J., Ben-David, E., & Burga, A. (2024). Selfish conflict underlies RNA-mediated parent-of-origin effects. Nature, 628(8006), 122–129. 10.1038/s41586-024-07155-z

Sasmita, A. O., Ong, E. C., Nazarenko, T., Mao, S., Komarek, L., Thalmann, M., Hantakova, V., Spieth, L., Berghoff, S. A., Barr, H. J., Hingerl, M., Börensen, F., Hirrlinger, J., Simons, M., Stevens, B., Depp, C., & Nave, K. A. (2025). Parental origin of transgene modulates amyloid-β plaque burden in the 5xFAD mouse model of Alzheimer’s disease. Neuron. 10.1016/j.neuron.2024.12.025

Srinivasan, R., Lu, T. Y., Chai, H., Xu, J., Huang, B. S., Golshani, P., Coppola, G., & Khakh, B. S. (2016). New Transgenic Mouse Lines for Selectively Targeting Astrocytes and Studying Calcium Signals in Astrocyte Processes In Situ and In Vivo. Neuron, 92(6), 1181–1195. 10.1016/j.neuron.2016.11.030

Winchenbach, J., Düking, T., Berghoff, S. A., Stumpf, S. K., Hülsmann, S., Nave, K. A., & Saher, G. (2016). Inducible targeting of CNS astrocytes in Aldh1l1-CreERT2 BAC transgenic mice. F1000Res, 5, 2934. 10.12688/f1000research.10509.1

Young, K. M., Mitsumori, T., Pringle, N., Grist, M., Kessaris, N., & Richardson, W. D. (2010). An Fgfr3-iCreER(T2) transgenic mouse line for studies of neural stem cells and astrocytes. Glia, 58(8), 943–953. 10.1002/glia.20976

Yu, X., Nagai, J., & Khakh, B. S. (2020). Improved tools to study astrocytes. Nat Rev Neurosci, 21(3), 121–138. 10.1038/s41583-020-0264-8

